# Differentiation of speech-induced artifacts from physiological high gamma activity in intracranial recordings

**DOI:** 10.1101/2021.04.26.441553

**Authors:** Alan Bush, Anna Chrabaszcz, Victoria Peterson, Varun Saravanan, Christina Dastolfo-Hromack, Witold J. Lipski, R. Mark Richardson

## Abstract

There is great interest in identifying the neurophysiological underpinnings of speech production. Deep brain stimulation (DBS) surgery is unique in that it allows intracranial recordings from both cortical and subcortical regions in patients who are awake and speaking. The quality of these recordings, however, may be affected to various degrees by mechanical forces resulting from speech itself. Here we describe the presence of speech-induced artifacts in local-field potential (LFP) recordings obtained from mapping electrodes, DBS leads, and cortical electrodes. In addition to expected physiological increases in high gamma (60-200 Hz) activity during speech production, time-frequency analysis in many channels revealed a narrowband gamma component that exhibited a pattern similar to that observed in the speech audio spectrogram. This component was present to different degrees in multiple types of neural recordings. We show that this component tracks the fundamental frequency of the participant’s voice, correlates with the power spectrum of speech and has coherence with the produced speech audio. A vibration sensor attached to the stereotactic frame recorded speech-induced vibrations with the same pattern observed in the LFPs. No corresponding component was identified in any neural channel during the listening epoch of a syllable repetition task. These observations demonstrate how speech-induced vibrations can create artifacts in the primary frequency band of interest. Identifying and accounting for these artifacts is crucial for establishing the validity and reproducibility of speech-related data obtained from intracranial recordings during DBS surgery.

## Introduction

Invasive brain recordings performed in awake patients undergoing clinically indicated neurosurgeries provide a unique opportunity to study speech production with better spatial and temporal precision than noninvasive neuroimaging methods. One of the advantages of intracranial recordings is a much higher signal-to-noise ratio (SNR) and larger measurable frequency range. This allows examination of frequency bands above 70 Hz that are typically unattainable with noninvasive methods due to volume conduction effects and a sharp attenuation in power at higher frequencies when passing the skull (Mukamel and Fried, 2012). Many assume that a higher SNR in the intracranial recordings makes them less susceptible to artifacts frequently observed in noninvasive recordings, such as electro-myographic artifacts due to eye, jaw, lip and tongue movement (Flinker et al., 2018; Lachaux et al., 2003; Llorens et al., 2011). Comprehensive quantitative examination of the quality of the signal and identification of potential sources of artifacts in intracranial recordings, however, have received very little attention. Several types of artifacts found in scalp EEG have been described in intracranial recordings, such as eye movement artifacts in fronto-anterior regions (Ball et al., 2009), and facial and mouth movement artifacts in electrodes close to temporal muscles (Otsubo et al., 2008). This suggestes that movement artifacts can contaminate intracranial LFP recordings acquired to study of the neural control of speech production.

Neural activity in the high gamma frequency band (60-200 Hz) tracks specific features of speech perception and production. For example, increased power in the high gamma frequency range has been observed in the superior temporal gyrus in response to auditory stimuli (Crone et al., 2001; Hamilton et al., 2018; Mesgarani et al., 2014), and in Broca’s and motor cortices during speech production (Edwards et al., 2010; Flinker et al., 2015; Mugler et al., 2018). High gamma activity recorded from the Rolandic cortex (pre and postcentral gyri) has been shown to track articulatory and/or acoustic features of speech sounds (Bouchard et al., 2013; Cheung et al., 2016; Chrabaszcz et al., 2019; Conant et al., 2018). Some recent advances have even made it possible to reconstruct speech from the brain’s activity in the high gamma band (Anumanchipalli et al., 2019; Martin et al., 2019), pointing at its potential utility for brain-computer interfaces (BCI) to develop speech prostheses. Contamination of the neural signal with audio acoustics therefore is a potential barrier to decoding the true electrophysiological correlates of speech production.

Given the impact that speech dysfunction can have in patients with movement disorders, and the fact that the role of subcortical regions in speech production are not well understood, we recently developed a strategy to simultaneously record from the cortex and the subcortical implantation target during DBS surgery. With the patient’s consent, it is possible to temporally place an electrocorticography (ECoG) electrode strip on the surface of the brain, a technique that has been used safely in over 500 patients (Panov et al., 2017; Sisterson et al., 2021). Here, we report the systematic identification and quantification of speech-induced artifacts in several types of intracranial electrophysiological recordings obtained using a speech production task performed during DBS implantation surgery. We show that this artifact is caused by mechanical vibrations induced by the produced speech, and that it can also be found in a ‘blank’ headstage pin not connected to any electrode. The results presented in this study encourage careful assessment of possible audio-induced artifacts in intracranial recordings obtained during speech production research. Additionally, we provide suggestions for data collection and analysis that may reduce the potential for false discoveries.

## Materials and Methods

### Participants

Participants were English-speaking patients with Parkinson’s disease (21M/8F, age: 65.6±7.1 years, duration of disease: 6.1±4.1 years) undergoing awake stereotactic neurosurgery for implantation of DBS electrodes in the subthalamic nucleus (STN). Dopaminergic medication was withdrawn the night before surgery. All procedures were approved by the University of Pittsburgh Institutional Review Board (IRB Protocol #PRO13110420) and all patients provided informed consent to participate in the study.

### Behavioral task

Participants performed a syllable repetition task intraoperatively. Subjects heard consonant-vowel (CV) syllable triplets through earphones (Etymotic ER-4 with ER38-14F Foam Eartips) and were instructed to repeat them. Triplets were presented at either low (~50dB SPL) or high (~70dB SPL) volume using BCI2000 (Schalk et al., 2004) presentation software. The absolute intensity was adjusted to each participant’s comfort level, keeping the difference between low and high-volume stimuli fixed at 25dB SPL. Participants were instructed to repeat the low-volume syllable triplets at normal conversation level and the high-volume triplets at increased volume, “as if speaking to someone across the room”. The syllables were composed of a combination of either of the 4 English consonants (/v/, /t/, /s/, /g/) and either of the 3 cardinal vowels (/i/, /a, /u/), resulting in 12 unique CV syllables. A total of 600 triplets were constructed which were divided among 5 presentation lists, with each session composed of 120 triplets. No syllable was repeated within a triplet. Syllable position and phoneme occurrence within a triplet and intensity level (low or high) within each session were balanced. The produced audio was recorded with an PRM1 Microphone (PreSonus Audio Electronics Inc., Baton Rouge, LA, USA) at 96 kHz using a Zoom-H6 portable audio recorder (Zoom Corp., Hauppauge, NY, USA). The average number of trials per session was 106 (range: 14-120). The duration of the utterances was of 1.3 ± 0.3 s (mean ± standard deviation pooled across subjects), with a median within-subject standard deviation of 0.14 s.

### Neural recordings

As part of the standard clinical DBS implantation procedure, functional mapping of the STN was performed with microelectrode recordings (MER), acquired with the Neuro-Omega recording system (Alpha-Omega Engineering, Nof HaGalil, Israel) using parylene-insulated tungsten microelectrodes (25 μm in diameter, 100 μm in length). LFPs were recorded from stainless steel macroelectrode rings (0.55 mm in diameter, 1.4 mm in length) located 3 mm above the tip of the microelectrode. The microelectrodes were oriented on the microtargeting drive system using three trajectories of a standard cross-shaped Ben-Gun array with 2 mm center-to-center spacing (Central, Posterior, Medial). The MER and LFP signals were recorded at a sampling rate of 44 kHz. The neural signal was referenced to the metal screw holding one of the guide cannulas used to carry the microelectrodes. Prior to initiating MER mapping, one or two subdural electrocorticography (ECoG) strips with 54 or 63 contacts each (platinum 1 mm disc contacts arranged in a 3×18 or 3×21 layout, with 3 mm center to center spacing, PMT Cortac Strips models 2110-54-001 and 2011-63-002) were placed through the standard burr hole.

ECoG strips were targeted to the left superior temporal gyrus (covering the ventral sensorimotor cortex) and left inferior frontal gyrus. Signals from ECoG electrodes and DBS leads were acquired at 30 kHz with a Grapevine Neural Interface Processor equipped with Micro2 Front Ends (Ripple LLC, Salt Lake City, UT, USA). Recordings were referenced to a sterile stainless-steel subdermal needle electrode placed in the scalp, approximately at the location of Cz in a standard EEG montage.

Two or three sessions of the syllable repetition task were performed by each subject when the microelectrode was positioned at different depths within the STN. The clinical setup evolved during the collection of this dataset between subjects 22 and 23, transitioning from a traditional stereotactic frame to the use of robotic stereotactic assistance (Faraji et al., 2020).

This change resulted in modification of the stereotactic frame’s mechanical coupling to the electrode micro-drive.

After the functional mapping phase patients were implanted with one of the DBS leads models shown in Table 1:

- Medtronic 3389: Platinum/Iridium, 4 cylindrical macroelectrodes, contact length 1.5 mm, 1.27 mm in diameter, 0.5 mm axial electrode spacing (Medtronic, Minneapolis, MN, USA).
- St. Jude 6172 short: Platinum/Iridium, directional lead with two central rings split in three contacts, 8 contacts total, length 1.5 mm, axial spacing 0.5 mm, lead diameter 1.27 mm, outer tubing material polycarbonate urethane (Abbott Neuromodulation, Austin, TX, USA).
- Boston Scientific DB-2202-45: Platinum/Iridium, directional lead with two central rings split in three contacts, 8 contacts total, length 1.5 mm, axial spacing 0.5 mm, lead diameter 1.3 mm, outer tubing material polyurethane (Boston Scientific Neuromodulation Corp, Valencia, CA, USA).

**Table 1.**
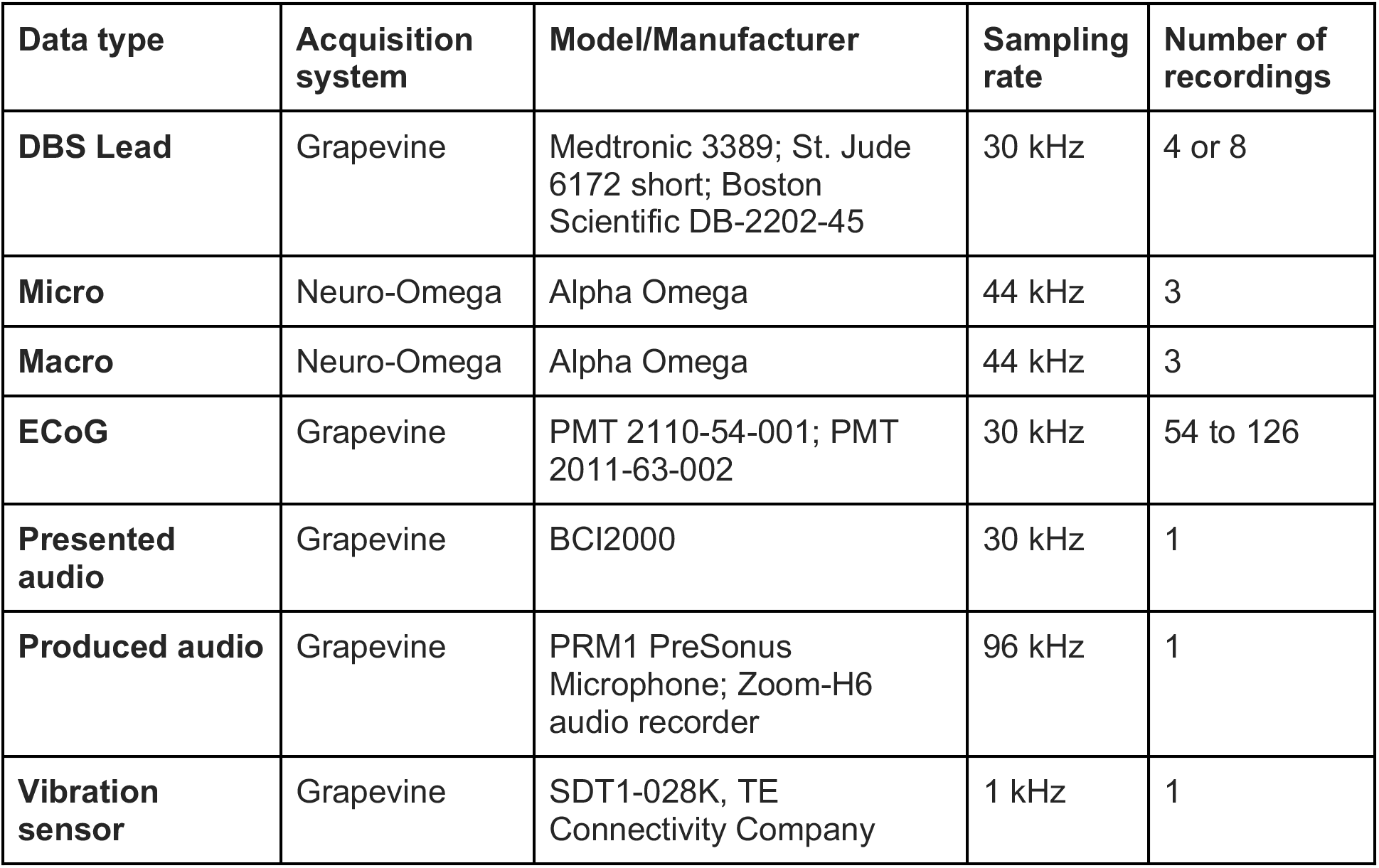
Acquisition system specifications for neural and physiological signals.

After the leads were successfully implanted, a final session of the speech task was performed by some participants, providing LFP data from DBS leads. A summary of recording types and the corresponding acquisition specifications is provided in Table 1. A schematic illustration of the intraoperative intracranial recording setup is shown in Figure 1.

**Figure 1:**
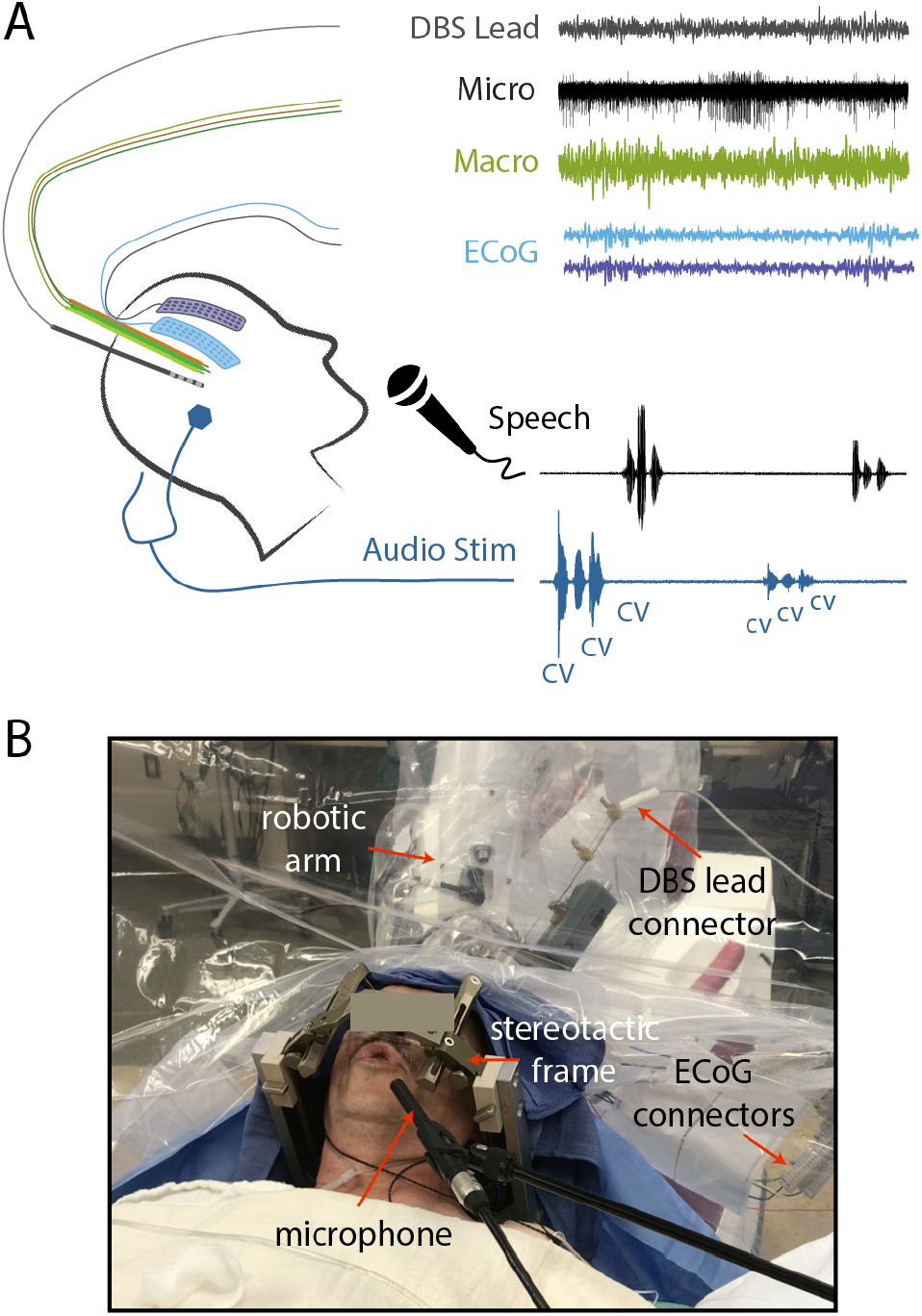
Intraoperative intracranial recording setup and speech task. **A.** Schematic representation of intracranial electrodes and syllable triplet repetition task. During the mapping phase of the DBS implantation surgery ECoG strips were temporarily placed through the burr hole, allowing simultaneous recording of cortical and subcortical LFPs and MER from the STN. Participants were instructed to repeat aloud CV syllable triples at a volume matching the auditory stimuli. **B.** Photograph of a participant performing the speech task.

### Vibration sensor

A shielded piezoelectric vibration sensor (model SDT1-028K, TE Connectivity Company) was fixed to the stereotactic frame using a sticky pad for the duration of one experiment. This sensor was selected for its flat transfer function at frequencies above 30 Hz. The signal from the sensor was captured without amplification as an analog input to the Grapevine system

### Electrode localization

DBS electrodes were localized using the Lead-DBS localization pipeline (Horn and Kühn, 2015). Briefly, a preoperative anatomical T1-weighted MRI scan was co-registered with a postoperative CT scan. The position of individual contacts was manually identified based on the electrode artifact present in the CT image and constrained by the geometry of the implanted DBS lead. This process rendered the coordinates for the leads in each subject’s native space. Based on the position of the lead and the known depth and tract along which the lead was implanted in each hemisphere, the positions of the micro and macro recordings from the functional mapping were calculated using custom Matlab scripts (github.com/Brain-Modulation-Lab/Lead_MER). The position of the ECoG strips was calculated from an intraoperative fluoroscopy image as described in (Randazzo et al., 2016). Briefly, the cortical surface was reconstructed from the preoperative MRI using the FreeSurfer image analysis suite (http://surfer.nmr.mgh.harvard.edu/). A model of the skull and the stereotactic frame was reconstructed from the intraoperative CT scan using OsiriX neuroimaging viewing tool (https://www.osirix-viewer.com/). The position of the frame’s tips on the skull and the implanted DBS leads were used as fiducial markers. The models of the pial surface, skull and fiducial markers were co-registered, manually rotated and scaled to align with the projection observed in the fluoroscopy image. Once aligned, the position of the electrodes in the ECoG strip was manually marked on the fluoroscopy image and the projection of those positions to the convex hull of the cortical surface was defined as the electrodes’ locations in the native brain space. The coordinates were then regularized based on the known layout of the contacts in the ECoG strip (github.com/Brain-Modulation-Lab/ECoG_localization). All coordinates were then transformed to the ICBM MNI152 Non-Linear Asymmetric 2009b space (Fonov et al., 2011) using the Symmetric Diffeomorphism algorithm implemented in the Advanced Normalization Tools (Avants et al., 2008). Anatomical labels were assigned to each electrode based on the HCP-MMP1 atlas (Glasser et al., 2016) for cortical electrodes, and the Morel (Niemann et al., 2000) and DISTAL (Ewert et al., 2018) atlases for subcortical electrodes.

### Phonetic coding

Phonetic coding of each participant’s produced speech was performed by a trained team of speech pathology students. Using a custom Matlab GUI (github.com/Brain-Modulation-Lab/SpeechCodingApp), onset and offset times, IPA transcription, accuracy and disorder characteristics of each produced phoneme were coded. Speech onsets were marked based on acoustic evidence of key speech features. Voice segments, such as vowels, were marked at the first visible glottal pulse, indicating the onset of vocal fold vibration. Unvoiced phonemes were marked based on the characteristic noise features for that phoneme. For example, the rapid increase of high frequency energy denoted the onset of plosive consonants. A broadband spectrogram was utilized to maximize visualization of key speech features. All coders completed a course in speech science to ensure knowledge of important speech features and were trained in coding procedures by a speech language pathologist.

### Electrophysiological data preprocessing

Data processing was performed using custom code based on the FieldTrip (Oostenveld et al., 2011) toolbox implemented in Matlab, available at (github.com/Brain-Modulation-Lab/bml). Recordings from the Grapevine, Neuro-Omega and Zoom-H6 systems were temporally aligned based on the stimulus and produced audio channels using a linear time-warping algorithm. Continuous alignment throughout the entire recording session was achieved with sub-millisecond precision. Data was low-pass filtered at 250 Hz using a 4^th^ order non-causal Butterworth filter, down-sampled to 1 kHz and stored as continuous recordings in FieldTrip datatype-raw objects in mat containers. This frequency range is well-suited for analyses in the canonical frequency bands normally used to explore cognitive functions. All annotations, including descriptions of each session (duration, type of subcortical recording, depth of the MER recordings), details of the electrodes (active time intervals, channel names, coordinates in native and MNI space, anatomical labels), phonetic coding at the phoneme, syllable and triplet level, and times of stimulus presentation were stored in annotation tables.

An automatic data cleaning procedure was used to remove segments of data with conspicuous high-power artifacts. First, a 1 Hz high-pass 5^th^ order non-causal Butterworth filter was applied to remove low frequency movement related artifacts. The power at frequencies in different canonical bands (3 Hz for δ, 6 Hz for θ, 10 Hz for α, 21 Hz for β, 45 Hz for γ_L_ and 160 Hz for γ_H_) was estimated by convolution with a Morlet wavelet with a width parameter of 9. For each frequency, the maximum power in 1-s time bins was calculated, log-transformed and the mean 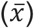 and standard deviations (σ) of the distribution were estimated using methods robust to outliers. A time bin was defined as artifactual if its maximal log-transformed power in any band exceeded 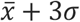 for that band. Note that this resulted in thresholds at least 10-fold higher than the mean power. Channels were entirely discarded if more than 50% of the time-bins were classified as containing artifacts. Blocks of channels sharing a head-stage connector were entirely discarded if more than 50% of those channels were defined as artifactual.

For each trial, 3 different behavioral epochs were defined: stimulus presentation or listening epoch - the 1.5-s long window during which syllable triplets were presented auditorily; speech production epoch - the variable time during which subjects repeated the syllable triplet; baseline epoch - a 500-ms time window centered between speech offset of one trial and stimulus onset of the following.

### Electrophysiology data analysis

#### Time-frequency plots

Time-frequency analyses for neural and audio data were performed using the Short Time Fourier Transform (STFT) method with a 100 ms Hanning window and a frequency step of 2 Hz, based on multiplication in the frequency domain as implemented by FieldTrip. Trials were aligned to speech onset and Z-scored relative to a 500-ms baseline epochs included in every trial. Frequencies up to 250 Hz were used for this analysis as that covers the canonical frequency bands normally used to explore cognitive functions.

#### Spectrogram correlation analysis

To index the degree of similarity between the audio spectrogram and the time-frequency spectrogram of a neural channel calculated by the STFT method, a correlation between these two matrices was calculated. This correlation was computed as the normalized sum of the element-by-element products of the two matrices.

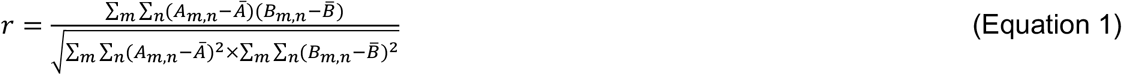

where A represents the audio spectrogram and B the time-frequency spectrogram of data at a given neural channel.

#### Coherence analysis

Phase relationship between the audio signal and the neural signal at a given channel was quantified using a metric of inter-trial phase consistency (ITPC) (Cohen, 2014). First, the audio and the neural signals were band-pass filtered between 70 and 240 Hz using a 5^th^ order non-causal Butterworth filter. This frequency range contains the fundamental frequency of the voice in humans and the narrowband component studied in this work. A notch filter was applied to remove line noise and its harmonics. For each individual epoch of interest (noted by the index *e*), and neural channel (*X*) a complex value *φ_e_* was calculated following

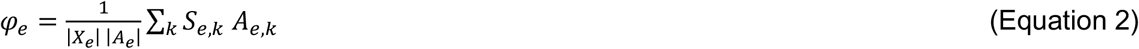

where *S_e_* = *X_e_* + *iH*(*X_e_*) is the analytic signal, *H*(*X_e_*) is the Hilbert transform of the neural data for the epoch *e*, and *A_e_* is the audio signal for that epoch. The sum is taken for every sample *k* within the epoch. |*X_e_*| and |*A_e_*| are the Euclidean norms of the neural and audio signals for epoch e. The absolute value and phase of *φ_e_* represents the degree of correlation and phase relationship between the neural and the audio channels for epoch *e*. If there is inter-trial phase consistency, the mean value of *φ* across trials (noted as 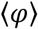) will be significantly different from 0. To quantify this, the ITPC index was defined as

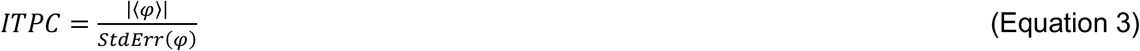

where the standard error of *φ* is defined as 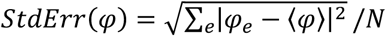, and the sum is taken over the *N* epochs considered for the neural channel of interest. Note that this metric was calculated independently for the speech production, listening and baseline epochs.

#### Significance threshold for coherence index

To define a threshold of significance for the coherence index, a Monte Carlo simulation was performed to obtain the distribution of coherence index under the null hypothesis of no consistent phase relationship between the neural and the audio channels. To this end, a random time jitter uniformly distributed between - 100 and 100 ms was applied to the neural data before calculating each *φ_e_* value. 22000 independent randomizations were calculated using data from all electrodes in the dataset. This analysis resulted in a significant threshold for the coherence index of 3.08 that corresponds to the 99.99 percentile of the distribution. Thus, a coherence index greater than 3.08 suggests significant correlation of inter-trial phase consistency between the audio and the neural signal.

#### Objective assessment of acoustic contamination

A recent study by Roussel et al. 2020, comparing intracranial recordings collected from human subjects during speech perception and production at five different research institutions found that spectrotemporal features of the recorded neural signal are highly correlated with those of the sound produced by the participants or played to participants through a loudspeaker. The method proposed by Roussel et al. for quantifying the extent of acoustic contamination in an electrophysiological recording was applied to the data using the open source toolbox developed by the authors (Roussel et al., 2020) (https://doi.org/10.5281/zenodo.3929296). Briefly, the method correlates the power of the neural data and audio across different frequency bins, thus creating a correlation matrix for every combination of frequencies of the two channels. Significantly higher correlation coefficients at matching frequencies of the two channels (i.e. on the diagonal of the matrix), compared to non-matching frequencies, is considered to be evidence of acoustic contamination. A permutation test is used to determine the significance threshold. The method was applied to data from individual electrodes, and the False Discovery Rate was adjusted according to (Benjamini and Hochberg, 1994).

#### Testing the spatial distribution of coherence over the brain

We used hierarchical bootstrapping (Saravanan et al., 2020) to test whether any particular brain region displayed an average coherence that was significantly different from the rest of the brain. Specifically, we built a null distribution of average coherence across the brain by calculating the mean coherence across the brain accounting for subjects as one hierarchical level but ignoring brain regions within subjects. We then calculated the average coherence for the 15 brain regions (as parcellated by the MMP1 atlas) that had coverage in 10 or more subjects, with 3 or more electrodes per subject. Each distribution was again built using subjects as the hierarchical level and electrodes were restricted to those within that particular brain region. Bootstrapping was performed weighing each subject by the number of electrodes present in that region. The distributions were then all compared to the null and the significance threshold was adjusted using an FDR correction for 15 comparisons. Resampling number (N_bootstrap_) was set to 10,000 for all bootstrap samples.

## Results

### Narrowband high gamma component in neural recordings during speech production

We averaged spectrograms time-locked to the speech onset across trials for each audio and neural channel. Representative examples of the averaged spectrograms for each recording type are provided in Figure 2. Figure 2A shows the spectrogram for the produced audio, in which the fundamental frequency (F0) of the participant’s voice (at around 120 Hz) and its first harmonic (at around 240 Hz) can be easily discerned as an increase in power at the corresponding frequencies. The 3 peaks of power in the spectrogram at different times correspond to the three produced syllables. A similar narrowband component occurring around the frequencies of the participants’ F0 also appeared in some electrodes during the speech production epoch. For example, this narrowband component can be readily observed in the time-frequency plot for one ECoG electrode shown in Figure 2B, but not for another ECoG electrode from the same subject shown in Figure 2C. Thus, while both electrodes show an increase in speech-related gamma activity, the increase in gamma activity in the electrode in Figure 2B is remarkably similar to the narrowband component in the audio spectrogram (Figure 2A), overlapping with it in frequency, time, and overall shape. Similar narrowband components in the high gamma frequency range were also identified in some of the LFP recordings extracted from the micro electrodes (Figure 2D), macro electrodes (Figure 2E) and from the DBS leads (Figure 2F). Thus, this narrowband speech-related component can be observed in different electrophysiological recordings collected simultaneously with different acquisition devices.

**Figure 2:**
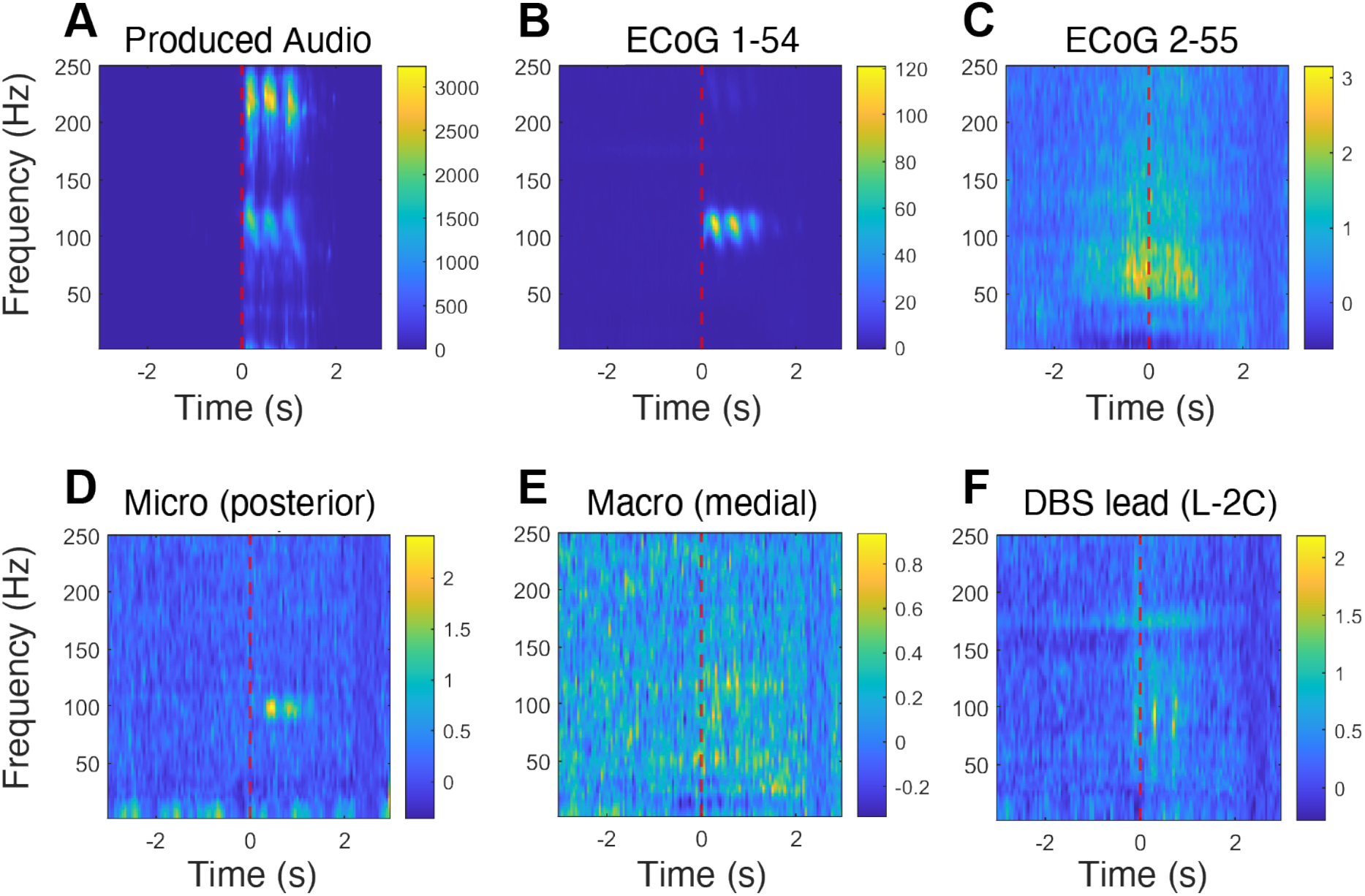
A speech-related narrowband component in the high gamma frequency range was observed across different types of neural recordings. Time-frequency plots for the audio and different neural channels from subject DBS3014. The red vertical dashed line represents speech onset time, which was used to time-lock the data across trials. **A.** Average spectrogram of the produced speech audio, z-scored to baseline. **B.** ECoG contact 1-54, showing a prominent narrowband high gamma component during the speech production epoch. **C**. ECoG contact 2-55, showing a broadband activation during speech production (note that this activity begins before speech onset). **D.** Spectrogram for LFP signal extracted from the posterior micro electrode targeting the left STN. **E.** Spectrogram for LFP signal from the medial macro ring targeting the left STN. **F.** Spectrogram for the left DBS lead contact.

### The narrowband high gamma component correlates with the fundamental frequency of the voice

Because of the striking similarity between the neural data and speech spectrogram, we set out to quantify the identified narrowband high gamma component and its relationship with the produced audio further, using different analytical approaches. First, we asked whether the observed overlap in the frequency range between the narrowband component and the produced speech audio was consistent across participants. To this end, we calculated the Welch power spectrum of the audio and neural data during the speech production epochs (Figure 3A), identified at which frequencies peak power within the F0 range (70-240 Hz) occurs in both types of spectra, and correlated the obtained frequency values. For each subject, we used data from single trials of the LFP signal extracted from one of the micro channels of the subcortical mapping electrodes, selected for having a prominent narrowband high gamma component.

**Figure 3:**
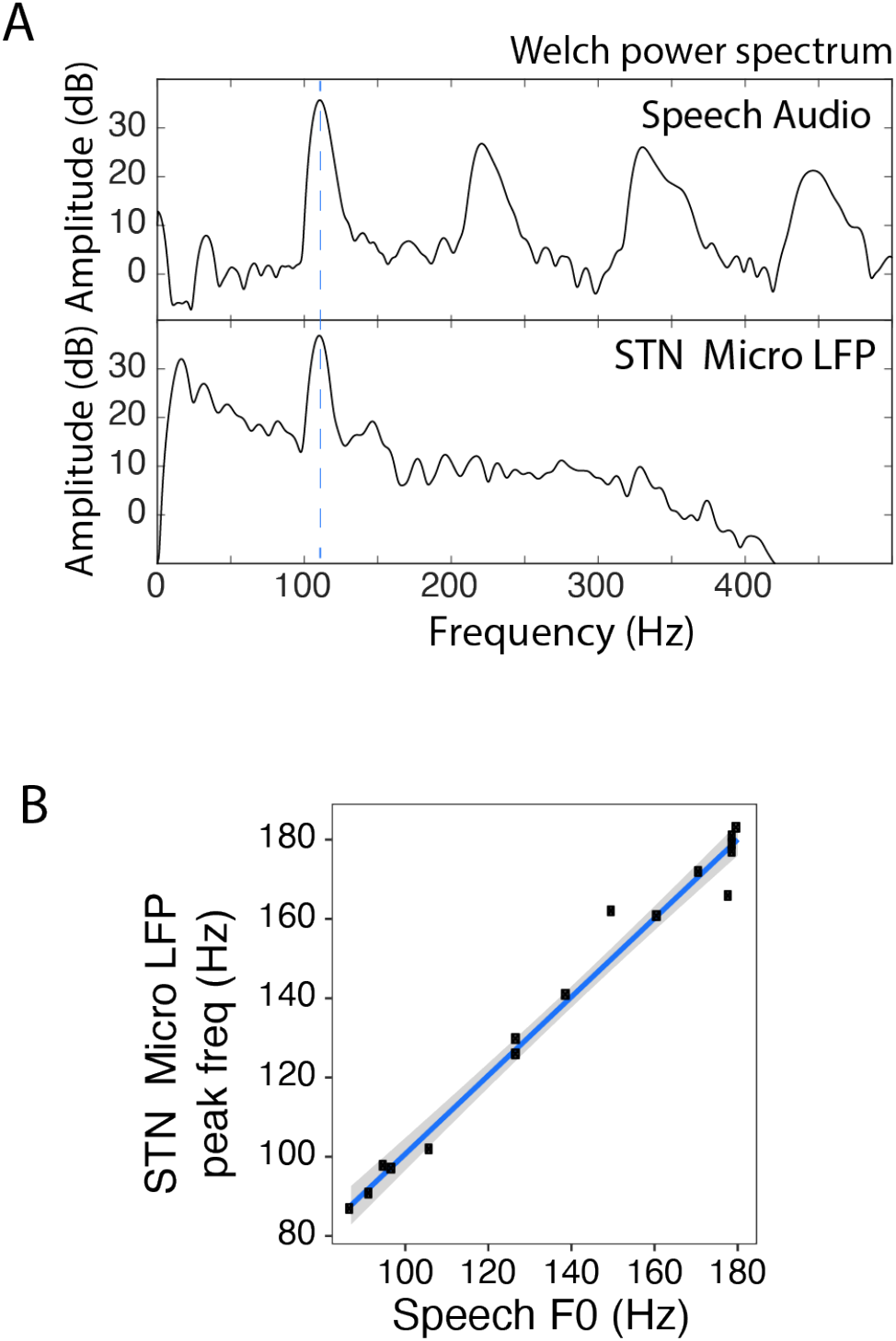
The frequency of peak power of the high gamma narrowband component correlates with the fundamental frequency of the voice across participants. **A.** Welch power spectrums for the produced speech audio (top) and the LFP signal extracted from the micro tip of the subcortical mapping electrode by low-pass filtering at 400 Hz (bottom). The vertical dashed blue line represents the frequency of peak power identified in each spectrum. **B.** Correlation of the frequency of peak power in the range of 70-200 Hz between the audio and neural data. Blue line represents the best linear fit to the data; gray ribbon represents the confidence interval of the fit.

As can be seen in Figure 3B, there is a strong correlation (Spearman’s p=0.98, p-value < 10^−6^, intercept = 1.4±5.1 Hz, slope = 0.99±0.4) between the fundamental frequency of the voice and the peak frequency of the narrowband component across participants. Furthermore, the relation not only is linear, but also has a slope not significantly different from 1, meaning that the frequency of peak high gamma power in the neural data corresponds to the fundamental frequency of the voice.

### Spectrogram cross-correlation between audio and neural data

To further characterize the distribution of this narrowband component across electrodes and recording sessions, we developed two different and complementary measures of similarity between the neural signal and the audio signal. The first metric consists of correlating the time-frequency spectrogram of a neural channel across times and frequencies with the spectrogram of the produced audio. A schematic representation of the method is shown in Figure 4A. For each subject, the correlation coefficient between the spectrogram of the produced audio and the spectrogram of neural recordings was calculated (Equation 1). We found widespread correlation between spectrograms obtained from intracranial recordings and spectrograms of the produced audio, reaching correlation coefficients of up to 0.8 (Figure 4B). The strength of correlations was variable across subjects, recording sessions, and electrodes. Also note that strong correlations were present in different types of intracranial recordings, albeit with a varying consistency.

**Figure 4:**
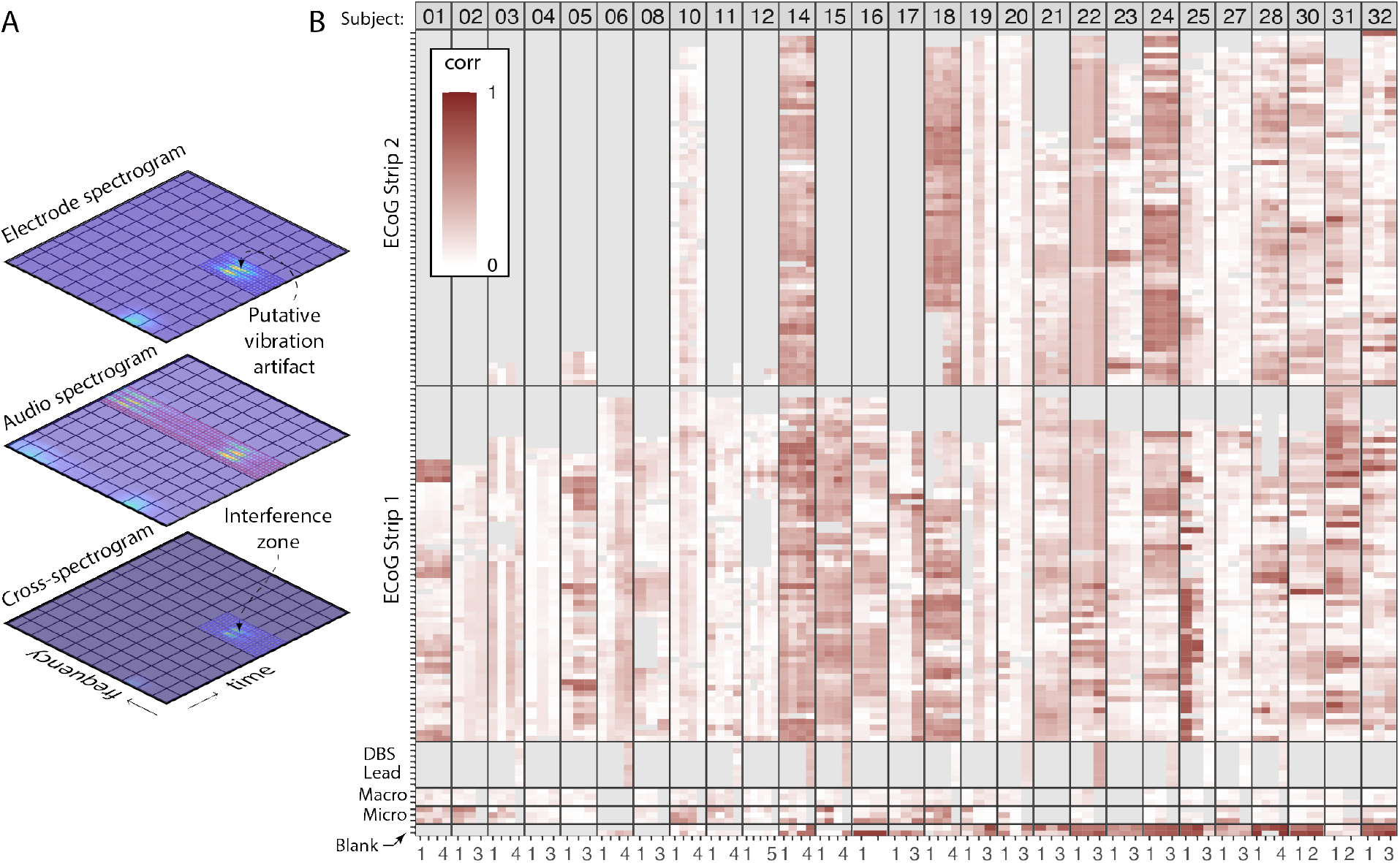
Spectrogram correlation between audio and neural data. **A.** A schematic representation of the spectrogram correlation approach. The electrode spectrogram (top) is multiplied by the audio spectrogram (middle) to give the cross-spectrogram (bottom). **B.** Correlation coefficients (Equation 1) for each electrode (represented on the y-axis), and each recording session (represented on the x-axis). Panels are defined by subject and electrode type. Tiles in gray correspond to electrodes not present in the montage or which were removed after the artifact rejection.

### Consistent phase relationship between audio and neural data

The method described above compares similarity of time-frequency resolved *power* between neural and audio data but does not take into account the phase information of the signals. Therefore, we developed a complementary metric to quantify the *phase* relationship between the neural data and the produced audio, that is, a measure of coherence or inter-trial phase consistency (ITPC) with the produced audio. This metric is based on dot products of the analytic signal of the neural channel and the produced audio, as schematized in Figure 5A. For each epoch, a complex value representing the magnitude of the correlation and the phase relationship of the neural signal with the audio is obtained (Equation 2). If the mean of these complex values is significantly different from zero, this serves as evidence of a consistent phase relationship between the neural data and the produced speech (Figure 5B). We used the absolute value of the mean of these complex values, normalized by their standard error, as an index of coherence (Equation 3). This method is computationally efficient and well suited for narrowband signals as the one we are characterizing (see details in Methods Section). We calculated a significance threshold using a Monte Carlo simulation in which we applied random time jitters before calculating the coherence index (see details in Methods Section).

**Figure 5:**
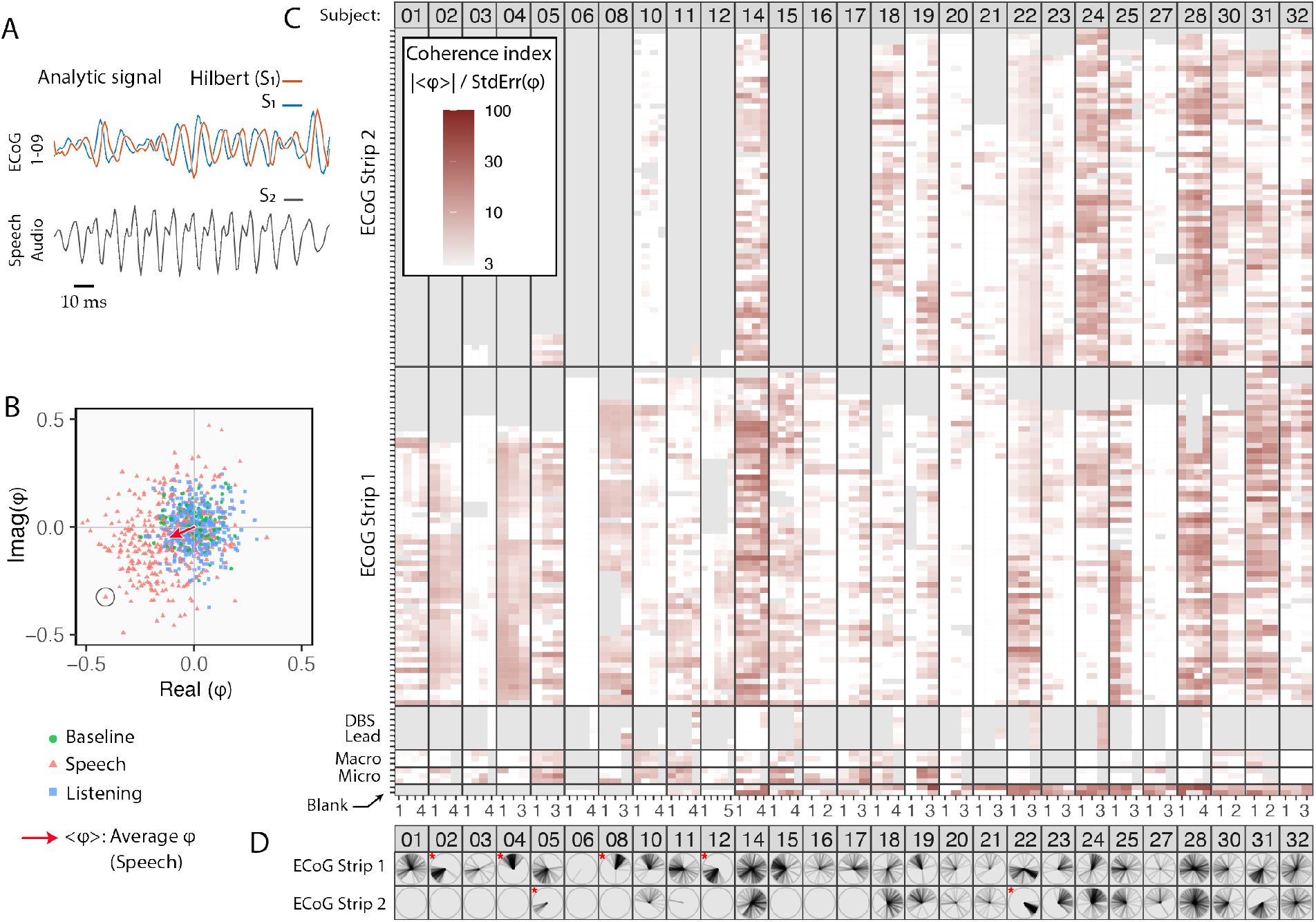
Inter-trial phase consistency (ITPC) reveals significant coherence of neural data with the produced speech audio. **A.** An ITPC index is derived from multiplying the band-pass filtered analytic neural signal (blue and orange lines for real and imaginary parts) with the produced speech signal (black line) using the internal product, resulting in a complex value *φ* for each trial. **B.** An example *φ* values plotted on the complex plane (real part of *φ* on the x axis and imaginary part on the y axis) for three behavioral epochs: baseline (green circles), listening (blue squares), and speech production (red triangles). The ITPC index is related to the average value of < *φ_e_* >, represented as a red arrow for the speech production epoch. For the baseline and listening epochs the < *φ_e_* > = 0, indicating no phase consistency between the neural and the audio data. The circled point in the bottom left quadrant of the plot corresponds to the traces shown in panel A. **C.** ITPC indices from the speech production epoch plotted for each electrode, recording session, and subject. Each row corresponds to an electrode; each column - to an individual session; panels are defined by subject and data type. Tiles in gray correspond to electrodes that were not present in the montage or which were removed after artifact rejection. White tiles represent electrodes that do not show significant coherence indices **D.** Phase consistency of the neural data with the produced audio across contacts of the ECoG strips. In each polar plot, the angle between the radial lines and the horizontal axis represents the phase relation with the produced audio of an ECoG electrode. Red asterisks mark ECoG strips with homogeneous coherence, defined as those with at least 90% of the electrodes’ phases within 90° of each other.

As observed in Figure 5C, there is widespread coherence across many subjects and all electrode types during the speech production epoch. Around 50% of the analyzed electrodes show significant coherence with the audio during the speech production epoch (Table 2). Importantly, no significant coherence with the speech audio was observed during the baseline or listening epochs (Table 2). Coherence of the neural signals with the stimulus audio in all behavioral epochs, including the listening epoch, was negligible (Table 2). This suggests that only the process of producing speech sounds was contaminating the neural signal.

**Table 2.** Percentage (± standard error) of electrodes showing significant similarity between neural data and the produced speech audio or stimulus audio, as established by the coherence method and the Roussel et al.’s method. The coherence method was run independently for the baseline, listening and speech production epochs. The Roussel et al. 2020 method used the entire duration of the session. Hierarchical bootstrapping was used to estimate the standard error.

| Audio | Produced speech audio |  |  |  | Stimulus audio |  |  |  |
| --- | --- | --- | --- | --- | --- | --- | --- | --- |
| Method | Coherence |  |  | Roussel | Coherence |  |  | Roussel |
| Epoch | baseline | listening | speech | all | baseline | listening | speech | all |
| **ECoG** | 0.2±0.1% | 0.2±0.1% | 52±6% | 54±5% | 0.4±0.3% | 0.2±0.1% | 1.4±0.7% | 2.2±0.5% |
| **Macro** | 0% | 0% | 42±8% | 25±5% | 0% | 0% | 0% | 4±2% |
| **Micro** | 0% | 0% | 56±7% | 96±7% | 0% | 0% | 0% | 0.6±0.8% |
| **DBS Lead** | 0% | 0% | 47±10% | 59±10% | 6±6% | 7±6% | 10±8% | 12±8% |
| **Blank** | 10±4% | 12±5% | 65±8% | 71±6% | 2±2% | 15±8 | 3±2% | 12±7% |

The distribution of the coherence indices across electrodes for each recording session can be classified as i) having no or little coherence between neural channels and the produced speech (e.g. subjects 06, 03, 21); ii) having homogenous coherence across electrodes, (e.g. subjects 02, 04); and iii) having heterogeneous coherence across electrodes (e.g. subjects 14, 28). While for the homogeneous case both the amplitude of the coherence (|<*φ*>|) and the phase relationship with the produced speech are similar across the neural electrodes, in the heterogeneous recording sessions these values may change substantially across electrodes (Figure 5D).

### The speech acoustic component can be detected in the neural data by different quantification methods

The spectrogram correlation index (Figure 4) and the coherence index (Figure 5) are highly correlated with each other, as can be observed in Figure 6A (Spearman’s p = 0.7, p < 1e-6). In applying the method proposed by Roussel et al. (2020), the power of the neural data was correlated with the audio across different frequency bins to create a correlation matrix (Figure 6B). High correlation coefficients on the diagonal of this matrix indicate acoustic contamination (see details in the Methods Section). Electrodes classified as contaminated based on this method are indicated by red triangles in Figure 6A. As can be observed, these points tend to have high spectrogram correlation index and coherence index above the significant threshold (Figure 6A). This result suggests that all three methods have similar sensitivity in identifying speech-related artifacts in our dataset.

**Figure 6:**
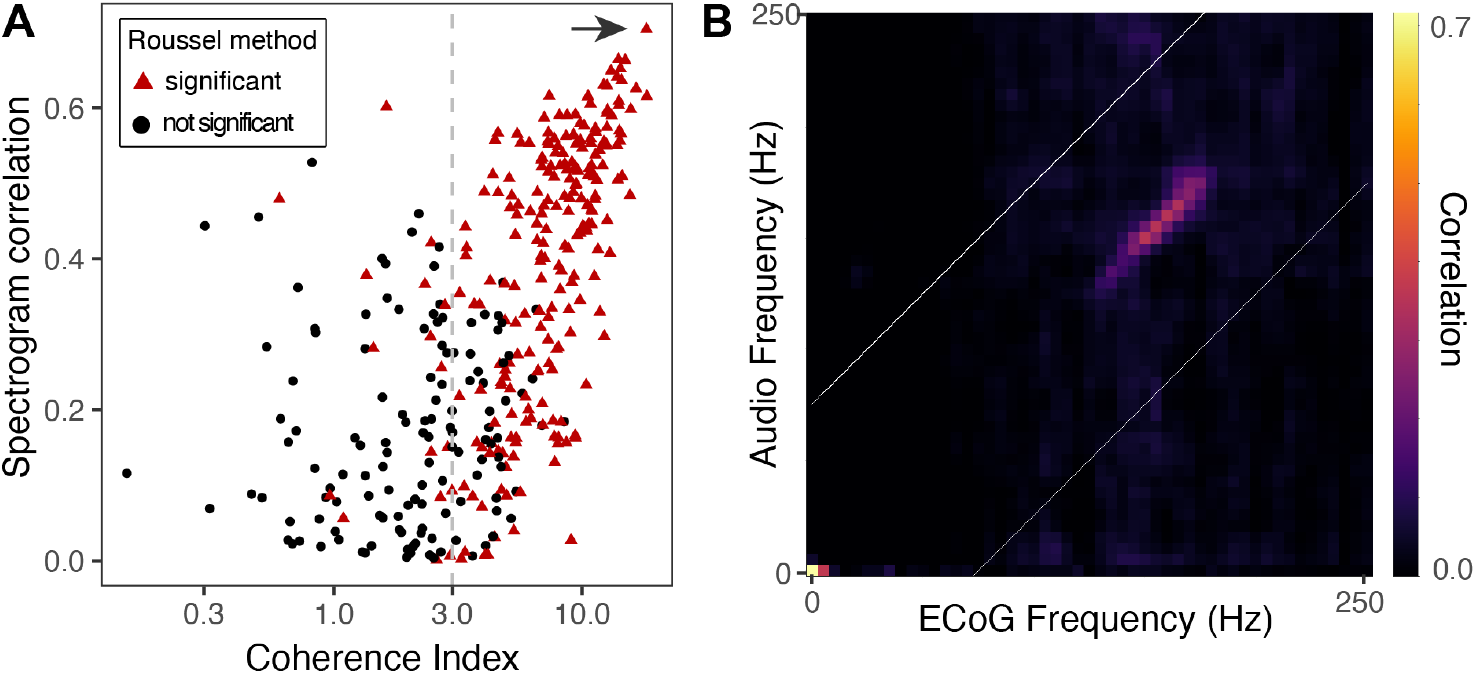
The spectrogram correlation index and the coherence index show strong correlation with each other and high consistency with the Roussel et al. method’s outcome. **A.** Relationship between the spectrogram correlation index and the coherence index for ECoG electrodes for one representative subject (DBS3024). Red triangles indicate electrodes with significant ‘acoustic contamination’ as assessed by the method developed by Roussel et al., 2020. **B.** Detection of acoustic contamination by the Roussel method is based on the cross-frequency correlation matrix, which indicates degrees of correlation of power across time for different frequencies of the neural and audio data. The shown matrix corresponds to ECoG electrode 2-31, indicated by an arrow in panel A.

### Coherence indices for cortical recording sites are independent of anatomic location

Next, we asked if there is any spatial structure of the coherence index within the cortex. To this end, we performed a hierarchical bootstrapping analysis accounting for the nested nature of the data (electrodes within anatomical regions within subjects) and found that none of the cortical regions (as defined by the Multi Modal Parcellation 1 in Glasser et al., 2016) show values of coherence significantly higher than the average distribution over the entire brain (see methods for detail). All cortical electrodes with their corresponding coherence values are plotted on a standard brain in Figure 7. Electrodes with high coherence with the produced audio do not cluster on any specific neuroanatomical region.

**Figure 7:**
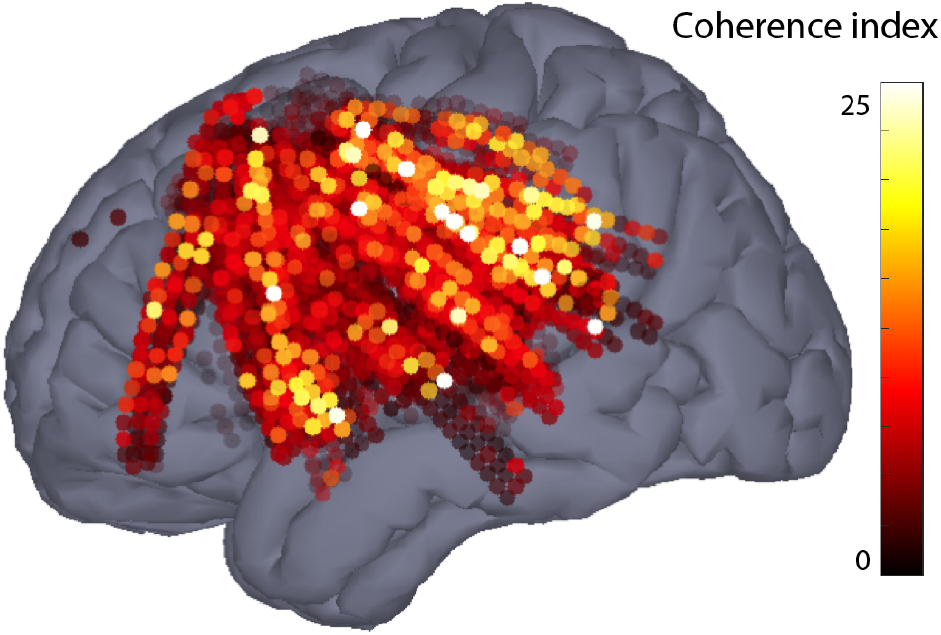
Increased Inter-trial Phase Consistency is not specific to any cortical region. Localizations of cortical ECoG electrodes for the entire subject cohort (n = 29) plotted in MNI space (MNI152 Nonlinear Asymmetric 2009b, (Fonov et al., 2011)). The color of the points represents the average coherence between neural and produced audio data for that electrode.

### Speech-related vibrations can be detected in non-neural data

The experiments described above demonstrated that some neural recordings show time-frequency patterns similar to the produced audio recordings. This suggests that there is contamination of the electrophysiological neural data with the speech audio signal. If this is the case, we expect to find the same kind of contamination in “blank” electrodes not in contact with the brain. Most of the ECoG strips used in our experiments contained 63 contacts laid out in a 3 x 21 arrangement (Figure 8A). These contacts were connected to the amplifier’s front-end through 4 cabrio-type connectors, each containing 16 pins. This resulted in the last pin of the 4^th^ cabrio connector (#64) not connected to any electrode on the brain. Instead, the wire connected to this pin runs the length of the cable and ends within the silicon rubber matrix of the ECoG strip. We recorded the signal from this “blank” pin as it provides a control for all non-neural sources of noise affecting the neural signal. Using the same time-frequency method as before, we analyzed the recorded signal from this blank ECoG headstage pin. As can be observed in Figure 8B, a narrowband component similar to that observed in some neural recordings is also present during speech production in the recordings from the blank headstage pin. It is interesting to note that the frequency of peak power recorded from this pin is slightly lower than the fundamental frequency of the voice (Figure 8C), although the component has the same timing and overall pattern (note the power increase around the first harmonic of the F0). As shown in the bottom row of the Figure 5C, labeled ‘Blank’, signals from the blank headstage pins show significant coherence with the produced speech audio in the same recording sessions that showed strong coherence with the produced speech audio in other electrode types (see also Figure 4B and Table 2).

**Figure 8:**
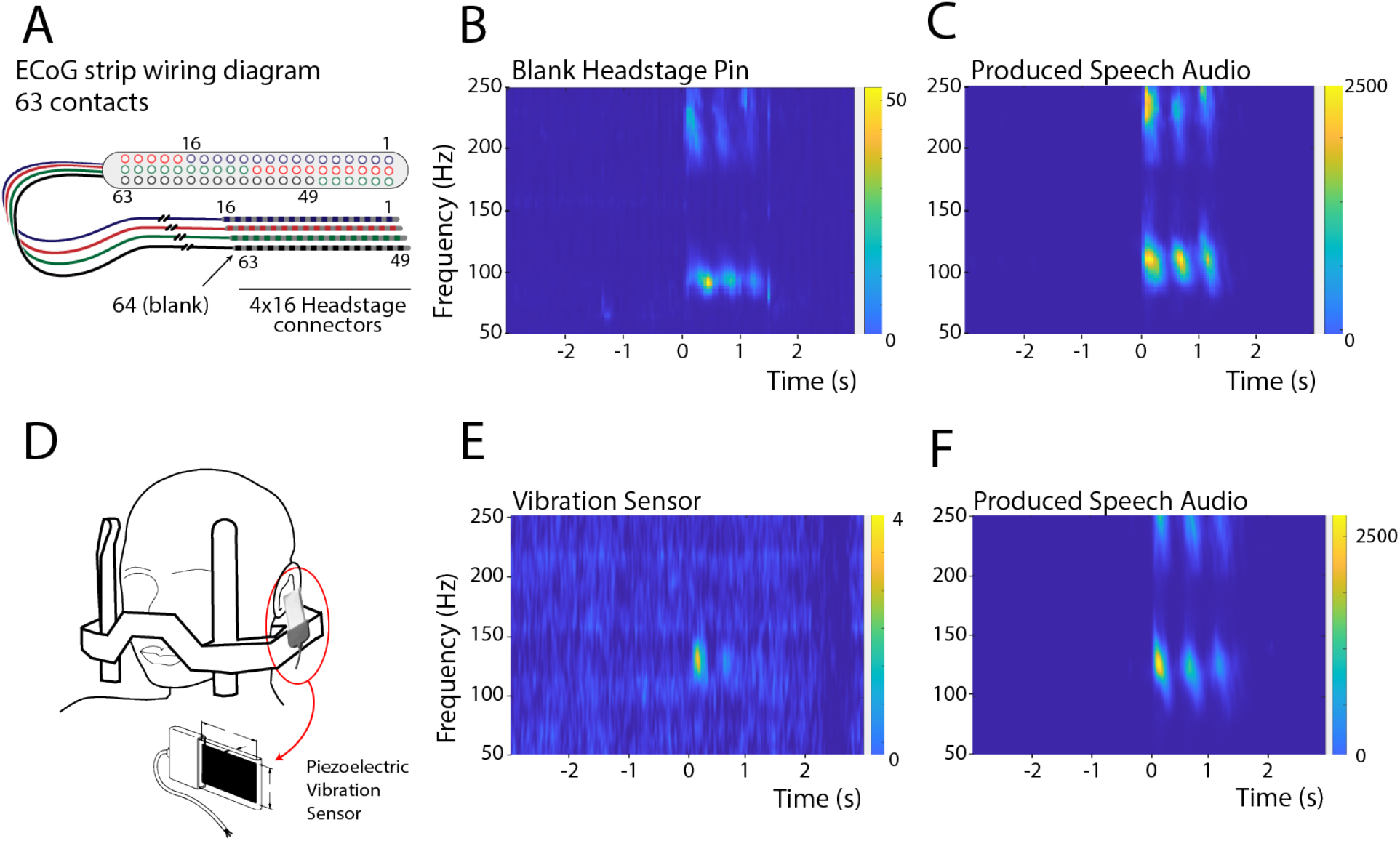
The narrowband component is present in non-neural recordings. **A.** A schematic of the ECoG electrode strip showing the electrode contact and connector layout. 4 cables of 16 wires are used for the 63 electrodes, resulting in one headstage pin not connected to any electrode on the brain (marked as contact #64). **B.** Time-frequency plot of the signal recorded from the blank headstage pin (subject DBS3020). **C.** Spectrogram of the produced speech audio for the same subject as in B. **D.** A schematic representation of the montage of the vibration sensor on the stereotactic frame. **E.** Time-frequency plot of the signal recorded from the vibration sensor (subject DBS3029). **F.** Spectrogram of the produced speech for the same subject as in E. In panels B-C and E-F, zero marks the onset of the speech production.

These results strongly suggest that the source of the observed narrowband component is not neural. Among possible sources of this component are speech-induced vibrations of the stereotactic frame, cables, connectors and/or acquisition chain. To address this question, we attached a piezoelectric vibration sensor to the stereotactic frame (Figure 8D) during data collection in one subject (subject DBS3029). The recorded signal from the piezoelectric sensor was analyzed in the same way as other types of recordings. A strong narrowband component was observed at the time corresponding to the production of the first syllable, in the same frequency range as the fundamental frequency of the voice (Fig. 8E). The attenuated power observed during the time of the production of the second and the third syllables might be due to the fact that in this particular participant speech intensity decreased across the produced syllable triplet, as seen in the audio spectrogram in Figure 8F.

## Discussion

We identified the presence of a narrowband high gamma component in the neural signals recorded from patients undergoing DBS implantation surgery that is consistent with a mechanically induced artifact. This component is widespread across many electrode types and was the most prominent feature in many electrode recordings. It occurs almost exclusively during cued speech production epochs and it has spectral and temporal characteristics strikingly similar to the produced speech audio. Indeed, a recent work shows the presence of acoustic contamination on ECoG recordings (Roussel et al., 2020). In the environment of DBS surgery, speech-induced vibrations conducted by the skull and stereotactic frame can impinge on the electrodes and/or signal acquisition chain affecting the recorded signal.

Several results support the interpretation that the observed narrowband high gamma component is an artifact due to speech-induced vibrations. First, the frequency of peak power of the narrowband component tracks the fundamental frequency of the voice across participants (Figure 3). Second, the narrowband high gamma component is almost exclusively present during the cued speech production epoch, but not the listening epoch (Table 2). Thirdly, the time-frequency resolved power from the neural recordings strongly correlates with the produced audio spectrogram (Figure 4). Fourth, significant inter-trial phase consistency between the produced audio and the neural data suggests similarities not only across the frequency domain, but also the temporal domain (Figure 5). Fifth, most of the recordings that we classified as contaminated in our analysis were also classified as having acoustic contamination by the recently proposed method in Roussel et al. (2020) (Figure 6 and Table 2). Sixth, there was no significant cortical localization of the ITPC index (Figure 7). Finally, the narrowband high gamma component was also detected in “blank” pins of the headstage connector not connected to any electrode (Figure 8B) and in the recording from a vibration sensor attached to the stereotactic frame (Figure 8E). Taken together, these results strongly suggest the presence of speech-induced vibration artifact.

Although we cannot rule out the presence of physiological neural activity with the exact same spectral and timing characteristics as the observed narrowband high gamma component, we favor the interpretation that the narrowband component identified in this work is mainly, if not completely, due to the vibration artifact. It is worth mentioning that broadband gamma activity can be detected in many electrodes (see example in Figure 2C), including some that also show the vibration artifact.

These results are in line with a recent report by Roussel et al. (2020), in which the authors show evidence for acoustic contamination of ECoG recordings. The authors analyze multiple-center datasets collected in epilepsy patients in an extra-operative setting, finding that both produced, and stimulus audio can affect ECoG recordings. Our work extends these findings, showing that the same kind of contamination can occur not only in the ECoG recordings, but can also affect other types of invasive neural recordings, such as those collected in the context of DBS implantation surgery. In addition, there are two key differences to note between our results and those of Roussel et al. First, we only observe contamination when the patient is speaking, but not during auditory stimulus presentation. This difference may be due to the use of headphones in our recording setup as opposed to the loudspeaker in the work by Roussel et al., who found that acoustically isolating the speaker reduced contamination of the neural signal. Second, in our recordings the affected signals are those corresponding to the F0 and to a lesser extent its harmonics. This could be explained by the fact that the stereotactic frame imposes some additional mechanical constraints, thus changing the way the vibration propagates.

Despite the fact that intracranial recordings in general are less prone to artifacts than extracranial recordings, it is recognized that the patient’s speech induces vibrations that affect MER. For example, a patent for a new micro-electrode design from AlphaOmega mentions that “*various mechanical noise and vibration, such as motor vibration, motion of the electrode within the tissue or voice of the subject, are detected by the electrode that acts essentially as microphone and is erroneously combined with the neural signal that is being recorded*” (Alpha Omega Technologies, 2020). Therefore, a common practice has been to analyze the recordings only when the patient is not speaking. The introduction of “microphonic free” microelectrodes with improved shielding that reduces the effects of vibrations on the recordings, allows acquiring single unit activity while the patient is speaking (Alpha Omega Technologies, 2020). Besides the clear clinical advantage, this has opened the possibility of studying single unit activity of DBS target structures during speech production. Although the quantifications of single-unit firing can be reliably performed with these electrodes (Lipski et al., 2017, 2018), in light of our results it is clear that mechanical vibrations affect other intracortical recordings including those obtained with macro contacts on the shaft of the micro electrodes, recordings from ECoG strips and recordings from the clinically implanted DBS leads.

Interestingly, neural signals with similar characteristics to the narrowband high gamma component described in this work have been reported in the literature. The Frequency Following Response (FFR) in the auditory brainstem is a potential with spectrotemporal features resembling the stimulus audio that reflects sustained neural ensemble activity phase-locked to periodic acoustic stimuli (Bidelman, 2018; Marsh and Worden, 1968; Marsh et al., 1970; Rose et al., 1966). This brainstem response to auditory stimuli can be recorded from scalp electrodes by averaging over hundreds of trials, a technique that has become a powerful diagnostic tool in audiology and neurology known as the Auditory Brainstem Response (Hall, 1992; Jewett et al., 1970). In recent years several works based on scalp EEG, sEEG and MEG have reported cortical FFRs (Behroozmand et al., 2016; Bidelman, 2018; Coffey et al., 2016).

In our data, there are two features that argue against the narrowband component being a true FFR. First, it only occurs during speech production and not during auditory stimulus presentation. Second, the narrowband artifact is not localized to the auditory cortex, or to any other cortical region (Figure 7). In light of the results presented in this work and similar results recently published by Roussel et al, it is clear that caution should be taken when interpreting cortical FFR, due to the fact that this signal is exactly what would be expected if there was an artifact (e.g., vibration at electrodes or connectors, electromagnetic induction by speakers, electrical crosstalk in the amplifier or connectors).

Other reported sources of artifacts affecting intracranial recordings, including artifacts due to eye blinking (Ball et al., 2009) and other facial muscle contraction (Otsubo et al., 2008), have spectral characteristics distinct from the narrowband component described in this work; they show broadband spectrums that do not match that of the produced audio and are not expected to track the fundamental frequency of the voice or correlate with the power of the speech audio across time.

Acknowledging the existence of this speech-induced vibration artifact is an important first step to avoid overinterpreting spurious features of the data. Many important questions related to high-level cognitive processes, including the neural control of speech production, can only be answered by acquiring recordings that are likely to be affected by the described artifact, but several steps can be taken to identify it. First, methods that correlate the audio signal with the neural data can be used to detect the presence of acoustic contaminations. Second, recording from open headstage pins provides a control for non-neural sources affecting the signal. Third, vibration sensors can detect mechanical vibrations along the recording system that might affect the signal. Furthermore, invasive recordings for BCI applications are likely to be affected by speech-induced vibrations, compromising the decoding performance. Therefore, it is necessary to develop source separation methods to remove speech-related artifacts from the neural data in order to reliably quantify underlying neural activity.

Identifying and mitigating artifacts in intracranial recordings from awake patients is fundamental to achieving reliable and reproducible results in the field of human systems neuroscience. This, in turn, will lead to an improved understanding of the neural physiology and pathophysiology of uniquely human cognitive abilities.

## Acknowledgments

We would like to thank all participants of this study for their time and effort. We would also like to thank Robert S. Turner for suggestions on data analysis. This work was funded by the National Institute of Health (BRAIN Initiative), through grants U01NS098969, U01NS117836 and R01NS110424 to R.M.R.

## Author Contributions

W.J.L. and R.M.R. designed the experiment and performed the recordings. A.B., A.C., C.D.-H. and W.J.L. reconstructed the electrode localizations and preprocessed the electrophysiological data. A.C. and C.D.-H. overviewed the phonetic coding. A.B., A.C., V.P., V.S., C.D.-H. analyzed data. A.B. and A.C. wrote the manuscript. All authors discussed the results and implications and commented on the manuscript at all stages. The authors declare no competing financial interests.

